# Coexistence holes characterize the assembly and disassembly of multispecies systems

**DOI:** 10.1101/2020.10.16.342824

**Authors:** Marco Tulio Angulo, Aaron Kelley, Luis Montejano, Chuliang Song, Serguei Saavedra

**Author notes:** These authors contributed equally (Chuliang Song, Serguei Saavedra).

## Abstract

A central goal of life science has been to understand the limits of species coexistence. However, we know surprisingly little about the structure of species coexistence below such limits, and how it affects the assembly and disassembly of ecological systems. Here we introduce a novel hypergraph-based formalism that fully captures the structure of coexistence in multispecies systems. Our formalism uncovers that, below its limits, coexistence in ecological systems has ubiquitous discontinuities that we call “coexistence holes.” These coexistence holes do not occur arbitrarily but tend to obey patterns that make them predictable. We provide direct evidence showing that the biotic and abiotic constraints of empirical systems produce an over-representation of coexistence holes. By highlighting discontinuities in the form of coexistence holes, our work provides a new platform to uncover the order and structure of the assembly and disassembly of ecological systems.

## Introduction

Ecological systems are the product of various assembly and disassembly processes (e.g., invasions and habitat fragmentation, respectively) that, together with the underlying ecological dynamics, determine which and how many species we observe^1–3^. Classic and recent studies have focused on characterizing the limits on the number of species that can coexist under given conditions via these two processes^4–8^. Yet, we know surprisingly little about the structure of species coexistence below those limits, and how it affects the assembly and disassembly processes^9^. Specifically, can these assembly and disassembly processes operate “smoothly” until reaching the limits of coexistence, as assumed in classical coexistence studies^10,11^? Or is it possible that they find “holes” where coexistence abruptly breaks before reaching the limits, causing discontinuities in both processes? Although these discontinuities—if they exist—reveal the internal constraints determining the structure of the assembly and disassembly of ecological systems^12,13^, we still lack a formalism to detect their occurrence and likelihood systematically.

Here, we address the gap above using a novel hypergraph-based formalism that fully captures coexistence in multispecies systems. Then, we use a new homology theory^14^ to analyze the structure of these hypergraphs, finding that discontinuities in the assembly and disassembly of ecological systems result in (topological) holes of different dimensions that we call “coexistence holes.” A coexistence hole occurs during the assembly when certain species collection is conceivable (i.e., it can be assembled from sub-collection that coexist), but it is not realized (i.e., it does not coexist). The opposite pattern gives rise to a coexistence hole during a disassembly process. We combine our formalism with a structuralist approach^9,12,15,16^ to build theoretical expectations about the emergence, likelihood, and structure of coexistence holes in ecological systems. We corroborate these expectations in five experimental microbial systems, finding additionally that the biotic and abiotic constraints in these empirical systems produce an over-representation of coexistence holes.

## Discontinuities in assembly and disassembly processes

### Coexistence holes

We consider ecological systems where individuals have been organized into *S* species (functional groups, taxa, or in any other meaningful organization). From the given species pool, each *species collection* either coexists or not. To study coexistence holes during the assembly process, we introduce the *assembly hypergraph H*. This hypergraph has the isolated species {1, 2, ⋯, *S*} as vertices, and it has one hyperedge *h* ⊆ {1, ⊆, *S*} for each different species collection that can coexist (Definition 2 in Supplementary Note S1). We illustrate this concept in the toy ecological system of *S* = 3 species of Fig. 1a. In this hypothetical ecological system, species survive in isolation and coexist when assembled in pairs, but species will not coexist when assembled together in the trio (Fig. 1b). The corresponding assembly hypergraph is *H* = [[1], [2], [3], [1, 2], [2, 3], [3, 1]]. By embedding (or realizing) the assembly hypergraph into an Euclidean space, discontinuities reveal themselves as holes in the hypergraph, which we call *assembly holes*. For example, embedding the assembly hypergraph *H* of the above toy ecological system into the plane reveals a one-dimensional assembly hole (Fig. 1c). The *dimension* of a hole is defined as the minimum dimension of its boundary. In our toy system, the hole’s boundary {[1, 2], [2, 3], [3, 1]} is one-dimensional, making the hole itself one-dimensional. A two-dimensional hole would appear, for example, in a hypergraph that looks like a tetrahedron with empty interior, and so on. Assembly holes occurs because, in all assembly processes to build certain species collection (e.g., to build [1, 2, 3]), coexistence abruptly breaks at the end (Fig. 1d).

**Figure 1:**
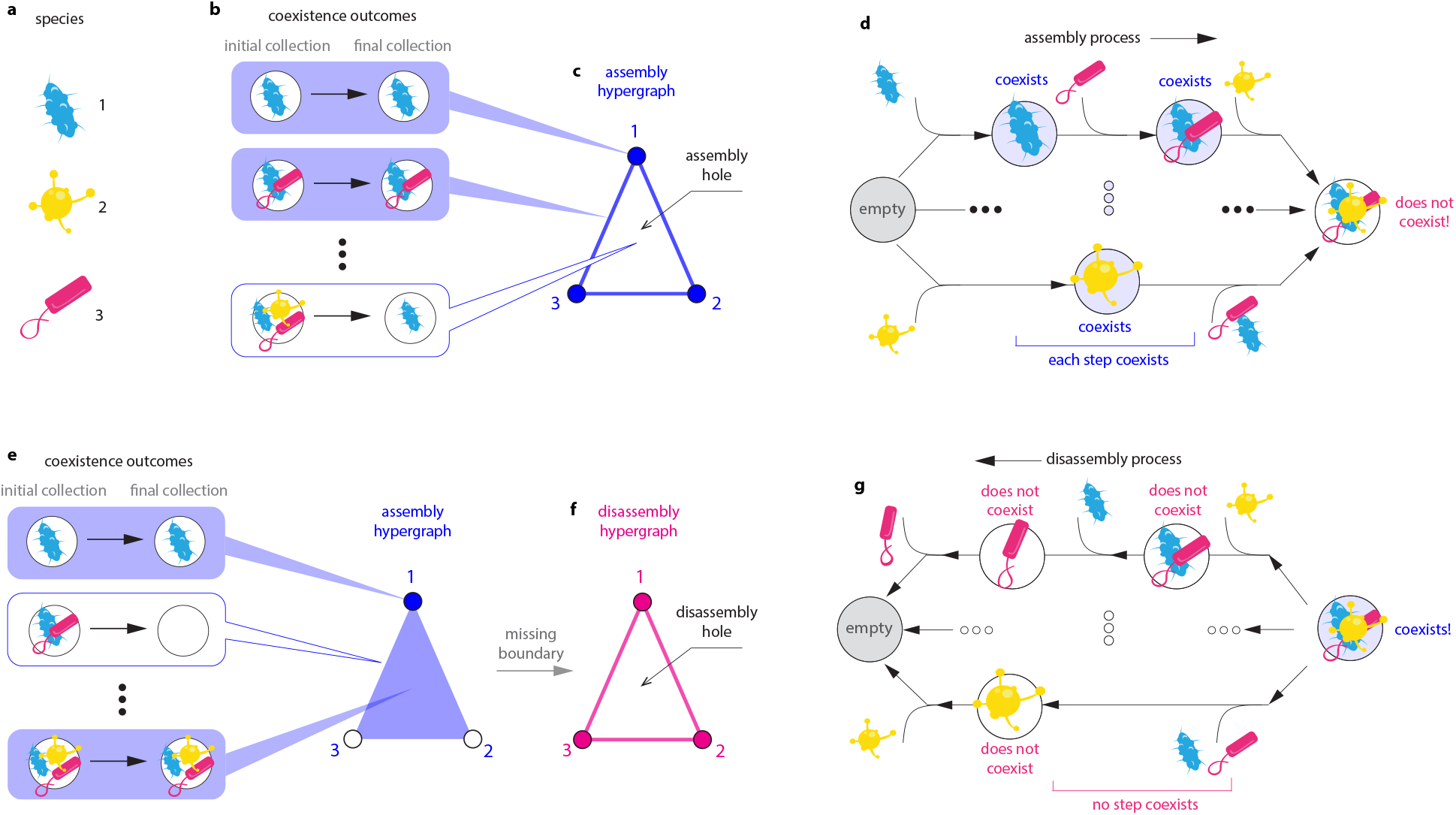
Coexistence holes characterize discontinuities in assembly and disassembly processes. **a.** A hypothetical pool of *S* = 3 species. **b.** When each of the 2*^S^* – 1 = 7 different species collection is assembled, it can either coexist (blue background) or not (white background). In this hypothetical example, species survive in isolation and coexist when assembled pairs. However, the three species cannot coexist when assembled together. The corresponding assembly hypergraph is *H* = [[1], [2], [3], [1, 2], [2, 3], [3, 1]]. **c.** When embedded into a two-dimensional space (i.e, plane), the assembly hypergraph reveals the assembly hole *h* = [1, 2, 3]. **d.** The assembly hole revels that coexistence abruptly brakes: in all assembly processes to obtain [1, 2, 3], all the intermediate species collections coexist, but in the final step coexistence does not occur. **e.** In these hypothetical coexistence outcomes, only species 1 survives in isolation, and coexistence is possible only if the species are assembled in a trio. The corresponding assembly hypergraph is *H* = [[1], [1, 2, 3]]. **f.** The associated disassembly hypergraph is *D*(*H*) = [[1], [2], [3], [1, 2], [2, 3], [3, 1]], calculated from the “missing boundary” of *H*. Each hyperedge of *D* is a sub-community that does not coexist, despite it was disassembled from the species collection [1, 2, 3] that coexists. When embedded into the plane, *D*(*H*) uncovers the disassembly hole [1, 2, 3]. **g** The disassembly hole reveals that coexistence abruptly brakes: despite [1, 2, 3] coexists, in all disassembly processes starting from [1, 2, 3], not a single intermediate species collection with more than one species coexists.

Moving on to study coexistence holes during the disassembly process, we introduce the *disassembly hypergraph D*. By definition, *D* contains all missing boundaries in the hyperedges of *H* (Definition 4 in Supplementary Note S1) Therefore, with the exception of zero-dimensional hyperedges (i.e., isolated species), each hyperedge of *D* is a species collection that does not coexist despite it was disassembled from a larger species collection that can coexist. To illustrate the disassembly hypergraph in an elementary case, consider the ecological system of *S* = 3 species with coexistence outcomes as in Fig. 1e. The assembly hypergraph for this hypothetical ecological system is *H* = [[1], [1, 2, 3]], see Fig. 1e. The corresponding disassembly hypergraph is *D* = [[1], [2], [3], [1, 2], [2, 3], [3, 1]]. By embedding the disassembly hypergraph into an Euclidean space, discontinuities in the coexistence reveal themselves as *disassembly holes*. For example, embedding the disassembly hypergraph of the above toy ecological system into the plane uncovers a one-dimensional disassembly hole (Fig. 1f). Disassembly holes occur because, in all disassembly processes starting at certain species collection (e.g., starting at [1, 2, 3]), coexistence abruptly breaks at their start (Fig. 1g).

### Uncovering assembly and disassembly holes via homology theory

Characterizing all discontinuities in the assembly and disassembly processes is equivalent to identifying all the coexistence holes of different dimensions in the assembly and disassembly hypergraphs. Identifying coexistence holes in systems with more than three species is challenging because it is likely that their assembly/disassembly hyper-graphs cannot be adequately embedded into a 2- or 3-dimensional space. To overcome this challenge, we constructed a novel *homology theory* allowing us to calculate the so-called *Betti numbers* of arbitrary hypergraphs (Methods and Supplementary Note S2). Here, the Betti numbers is the vector [*β*_0_, *β*_1_, ⋯] where *β_k_*, *k* ≥ 1, counts the number of k-dimensional holes in the hypergraph, and *β*_0_ counts the number of connected components (i.e., 0-dimensional holes). For example, the Betti numbers for the assembly hypergraph of Fig. 2a are [2, 3, 0, 0, ⊆], implying it has 2 connected components and 3 one-dimensional assembly holes. The Betti numbers for the disassembly hypergraph of Fig. 2b are [4, 1, 1, 0, ⋯], implying (among other things) it has 1 two-dimensional disassembly hole (made of the species collection [5, 6, 7, 8, 9], which is not obvious from the embedding of the hypergraph into the plane). By calculating Betti numbers, we can distill the “essential form” of assembly/disassembly hypergraphs into their (homotopic) assembly/disassembly *skeletons*—namely, a “minimal” hypergraph with the same Betti numbers (see Fig. 2c-d).

**Figure 2:**
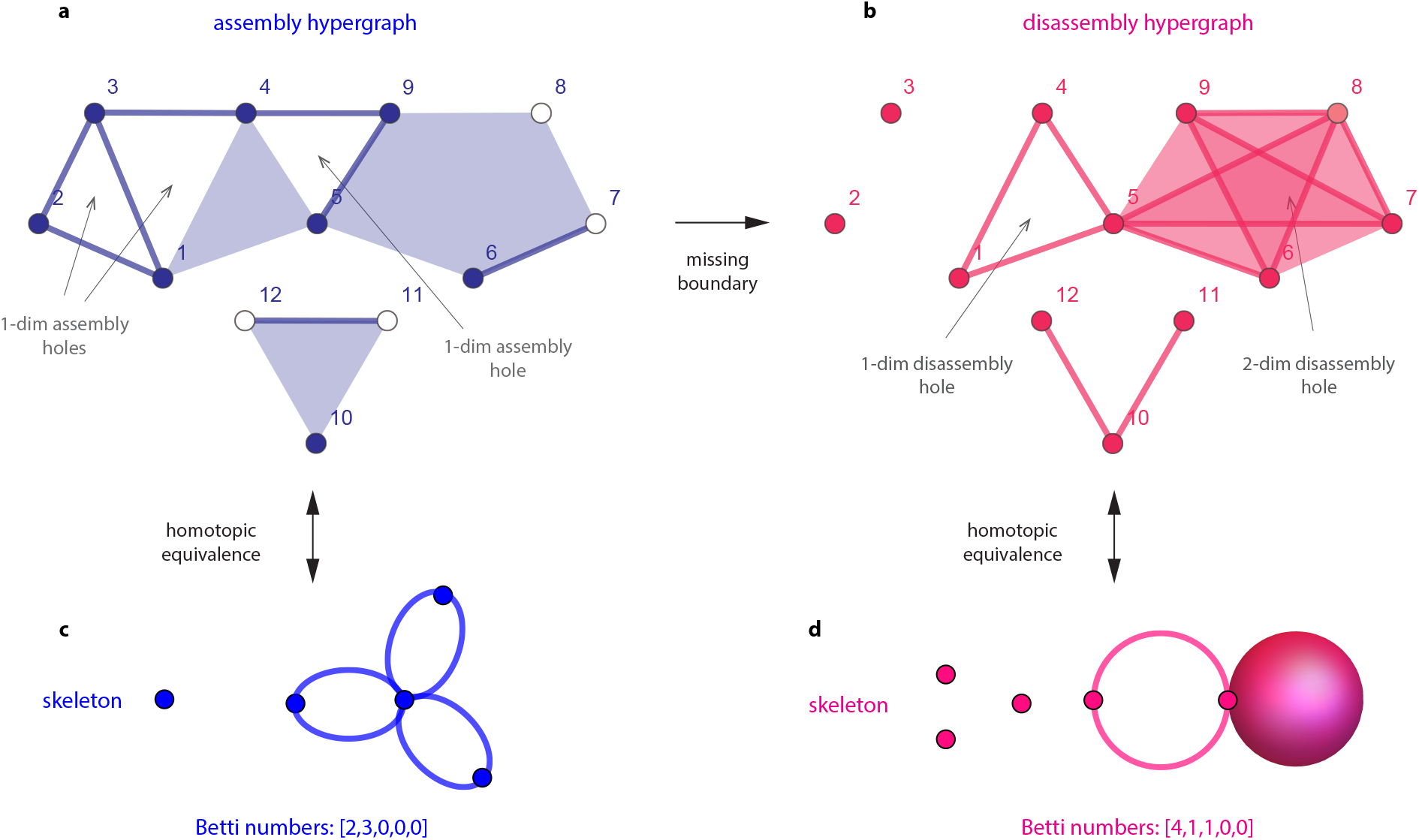
Homology reveals the essential structure of assembly and disassembly hypergraphs in terms of their skeletons. **a.** An assembly hypergraph for an hypothetical system of *S* = 12 species. While coexistence in this example is limited to 5 species, there exists additional structure within this limit. In particular, this assembly hypergraph contains 2 connected components (i.e., [10, 11, 12] and the rest) and 3 one-dimensional assembly holes. **b.** Homotopic skeleton of the assembly hypergraph of panel a. The skeleton is quantified by its Betti numbers. **c.** The disassembly hypergraph corresponding to the assembly hypergraph of panel a. The disassembly hypergraph contains 4 connected components, 1 one-dimensional hole, and 1 two-dimensional hole. Since this hypergraph is embedded in the plane, its two dimensional hole is not obvious. **d.** Homotopic skeleton of the disassembly hypergraph of panel c. The skeleton is quantified by its Betti numbers.

## Assembly and disassembly holes in theoretical ecological systems

### Emergence of coexistence holes

To study how discontinuities (i.e., assembly and disassembly holes) emerge in ecological systems, we combine our formalism with a structuralist approach^12,16^ that leverages on the mathematical tractability of the Lotka-Volterra (LV) model (Methods and Supplementary Note S3). Despite its simplicity, the LV model has successfully explained and predicted the ecological dynamics of diverse systems^16–22^. In our structuralist approach, we assume that biotic conditions characterize the internal constraints defining the structure of ecological systems, while abiotic conditions can change freely giving rise to the potential diversity within such internal constraints^15,23^. In the idealized governing laws of the LV model, biotic conditions are phenomenologically captured by the 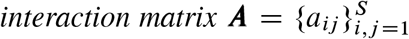, with *a_ij_* representing the effect of species *j* on the per-capita growth rate of species *i*. In turn, abiotic conditions are phenomenologically captured by the 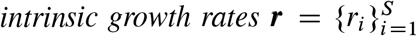, with *r_i_* representing how species *i* grows in isolation under given abiotic conditions. Here, free abiotic conditions imply that ***r*** can freely change over the unit sphere ∥***r***∥_2_ = 1^15,16^.

The consequence of the structuralist approach is that the interaction matrix ***A*** determines the possible assembly and disassembly skeletons that an ecological system can adopt, while the intrinsic growth rates ***r*** determine the observed skeleton out of the possible ones. For example, consider the hypothetical ecological system of *S* = 6 species interacting as described by the interaction matrix ***A*** of Fig. 3a. From this interaction matrix, a fixed and finite set of possible assembly and disassembly skeletons emerge (Fig. 3b). Each one of these skeletons is compatible with a specific range of directions of ***r***. This implies that each assembly and disassembly skeleton will be selected from the possible pool of skeletons with different probability. In our example, we have 6 possible assembly skeletons and 12 possible disassembly skeletons (Fig. 3c). The possible assembly and disassembly skeletons contain 3 assembly holes and 3 disassembly holes (Fig. 3d). The probability distribution over assembly skeletons is strongly concentrated on the skeleton [1, 0, 0, 0, 0], whereas the probability distribution over disassembly skeletons tends to be more uniform. More “complex” ecological systems—obtained by increasing the strength or number of interspecific interactions^24^—can adopt a larger number of assembly and disassembly skeletons (Supplementary Note S4.1 and Supplementary Fig. S4)

**Figure 3:**
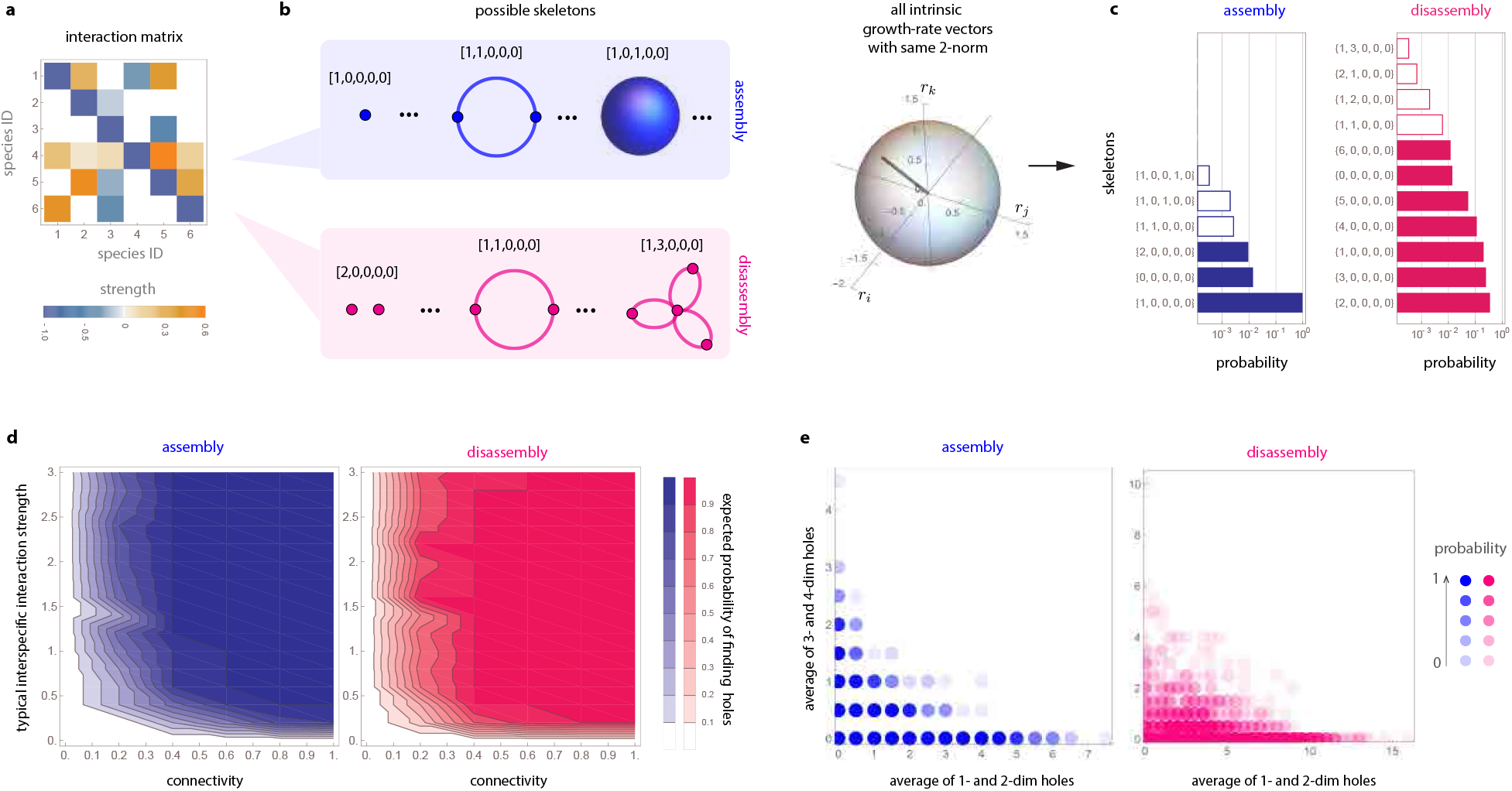
Assembly and disassembly holes in theoretical ecosystems. **a.** An hypothetical interaction matrix **A** describing the interactions between *S* = 6 species (see values in Supplementary Eq. (S6)). **b.** Possible assembly and disassembly skeletons generated by the interaction matrix of panel a. **c.** By choosing **r** uniformly at random over the unit sphere, some assembly/disassembly skeletons are more likely to occur. Here, solid bars denote skeletons without holes, and hollow bars denote skeletons with at least one hole. **d.** Expected probability of finding at least one assembly (blue) or disassembly (pink) hole in the ensemble of random LV models of *S* = 8 species. **e.** Not all possible assembly/disassembly skeletons are possible in ensemble of random LV models. Namely, note that skeletons that simultaneously have a large number of low-dimensional holes (i.e., 1- or 2-dimensional holes) and high-dimensional holes (i.e., 3- and 4-dimensional holes)—which occupy the top-right corner of the panels—are very unlikely.

### Simple dynamics generate assembly and disassembly holes

The hypothetical ecological system of Fig. 3a shows that discontinuities in the form of assembly and disassembly holes emerge even in simple population dynamics like the LV model, without sophisticated mechanisms such as complex functional responses^25^, higher-order interactions^26^, or historical contingency^1^. To investigate how general this conclusion is, we systematically analyzed an ensemble of systems governed by LV dynamics with random interaction matrices ***A*** (Supplementary Note 4.3). The ensemble is characterized by two parameters of the interaction matrix: its the typical interspecific interaction strength *σ_A_* > 0, and its typical connectance *C_A_* ∈ [0, 1]. We found that, as soon as the interaction matrix exceeds a small complexity threshold measured by *σ_A_C_A_*, the presence of assembly and disassembly holes is unavoidable (Fig. 3d). This result implies that discontinuities are the norm rather than the exception in the assembly and disassembly of random (unstructured) LV systems.

We also found that coexistence holes in unstructured ecological systems obey two general patterns that makes them predictable. First, knowledge of the disassembly skeleton strongly determines which assembly skeleton the system can adopt, but not vice versa (Supplementary Fig. S6). That is, there is an asymmetry in the information contained in the assembly and disassembly skeletons (Supplementary Note S4.5). Second, low-dimensional and high-dimensional assembly/disassembly holes are unlikely to co-occur. That is, when the number of low-dimensional (i.e., 1- or 2-dimensional) and high-dimensional (i.e., 3- or 4 dimensional) holes are used as coordinates in a plane, the assembly and disassembly skeletons generated by random LV models avoid the upper-right corner (Fig. 3e). This pattern arises because the likelihood of observing assembly or disassembly holes depends on their dimensions and the particular “complexity” of the interaction matrix (Supplementary Note S4.4). For example, high-dimensional assembly holes are more likely at low interspecific interactions, whereas low-dimensional holes are more likely at large interspecific interactions (Supplementary Fig. S5). Note that, mathematically, it is possible to conceive hypergraphs that simultaneously have many low- and high-dimensional holes. Therefore, the absence of such skeletons shows how the internal constraints of ecological systems shape the likelihood to observe certain discontinuities.

## Coexistence holes in empirical ecological systems

### Description of the empirical ecological systems

To study if coexistence holes occur in empirical ecological systems, we analyzed five experimental microbial communities: a system of *S* = 4 protozoa built by Vandermeer^21^, a system of *S* = 8 soil bacterial species built by Friedman et al.^27^, a system of *S* = 11 human gut bacterial species built by Stein et al.^19^, a system of *S* = 12 human gut bacterial species built by Venturelli et al.^18^, and an in-vivo system of *S* = 14 gut bacterial species built for the MDSine project^28^ (see details in Supplementary Note S5). In each of these studies, a LV model was parameterized and systematically validated using high-resolution experimental data, resulting in empirical estimations of the parameters (***A***, ***r***).

### Empirical species interactions generate coexistence holes

We find that all five empirical interaction matrices can generate assembly and disassembly holes (Fig. 4a-e). Indeed, while the numbers of possible assembly and disassembly skeletons differ across ecological systems, most of these skeletons contain assembly and disassembly holes. In the four ecological systems with larger species pools, the empirical interaction matrices generate between 1.2 (Friedman) and 12.0 (MDSine) times more disassembly skeletons than assembly skeletons (confirming our theoretical expectations about differences between assembly and disassembly skeletons). Importantly, these interaction matrices with larger species collections tend to have a larger proportion of skeletons with coexistence holes (i.e., with discontinuities). We also confirm our expectations that low- and high-dimensional assembly/disassembly holes are unlikely to co-occur (grey points in Fig. 4f-j), and that disassembly skeletons strongly influence assembly skeletons in all ecological systems, but not vice versa (Supplementary Fig. S7). For example, in the Stein system, disassembly reduces the uncertainty of assembly in about 45%, while assembly reduces the uncertainty of disassembly in about 20%. In the MDSine system, disassembly reduces uncertainty of assembly in 35%, while assembly reduces the uncertainty of disassembly in less than 5%.

**Figure 4:**
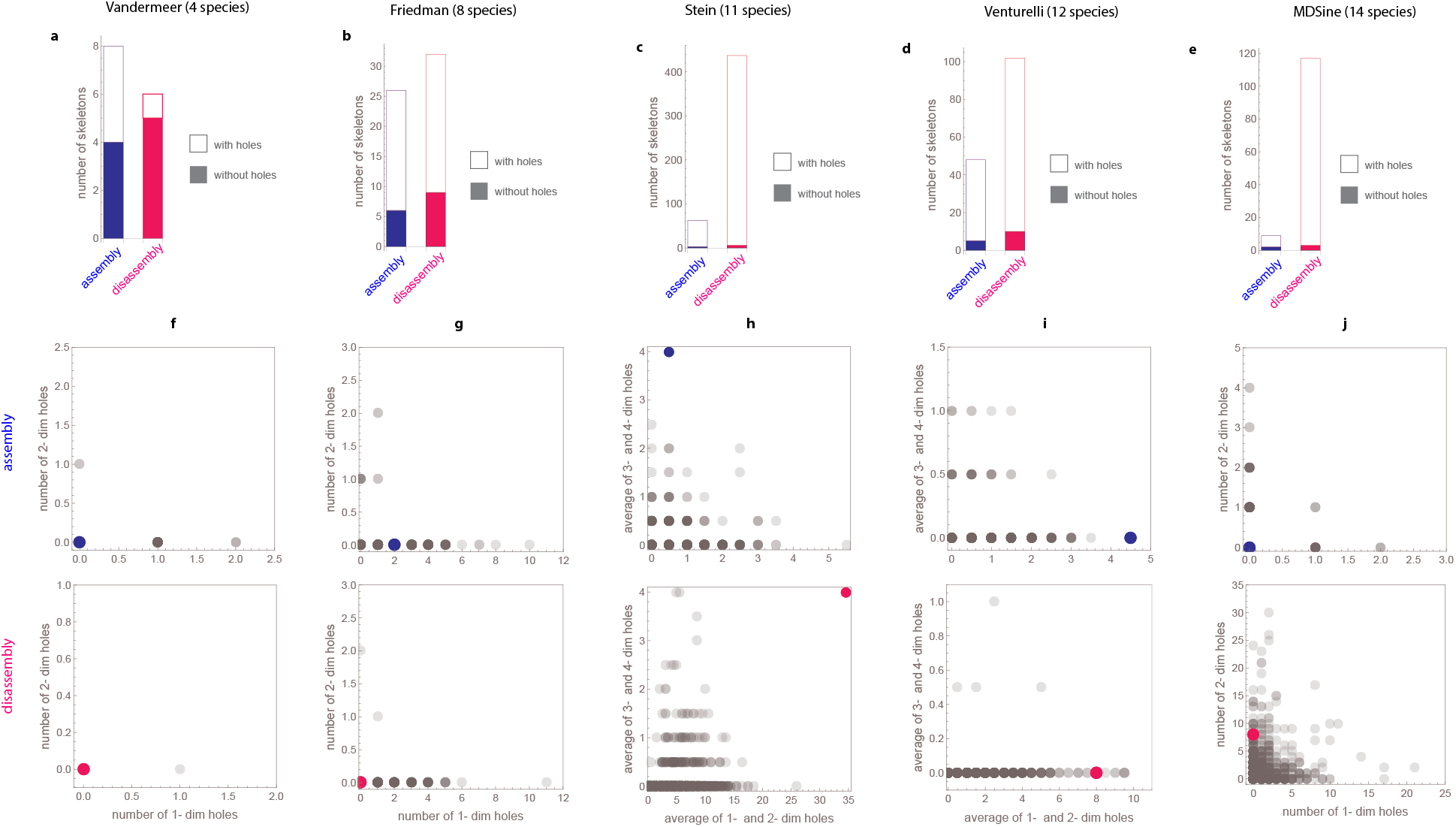
Empirical species interactions generate assembly and disassembly holes. Results are for each empirical interaction matrix and 3,000 skeletons generated by choosing the intrinsic growth-rate vector uniformly at random over the unit sphere. bf a. to e. Number of assembly and disassembly skeletons generated by each empirical interaction matrix. **f. to j.** Grey dots show the number of low- and high-dimensional holes observed in the 3,000 random skeletons obtained by using the empirical interaction matrices (strength of color is proportional to the probability of occurrence). Color dots show the empirical skeletons obtained by the empirical interaction matrix and the empirical intrinsic growth-rate vector.

### Empirical ecological systems are enriched with coexistence holes

Next, we focus on the five empirical assembly and disassembly hypergraphs obtained from the experimentally parameterized ***A*** matrices and ***r*** vectors (Fig. 5a-e). Our analysis shows that coexistence holes are over-represented in the assembly and disassembly skeletons of all five empirical systems compared to skeletons obtained by randomly changing the abiotic conditions. The Vandermeer and Friedman systems are enriched with 0-dimensional disassembly holes (*p* = 0.057 and *p* = 0.0043, respectively, Fig. 5f-g). Interestingly, the assembly hypergraph of these two systems is a simplicial complex—each hyperedge contains all its boundaries—suggesting a simple assembly principle. The Stein system is enriched with 3-dimensional assembly holes and 2-dimensional disassembly holes (*p* = 0.00075 and *p* < 0.00033, respectively, Fig. 5h). Remarkably, this empirical system contains 64 two-dimensional disassembly holes. The Venturelli system is enriched with 2-dimensional assembly holes and 1-dimensional disassembly holes (*p* < 0.00033 and p = 0.00025, respectively, Fig. 5i). For the MDSine system, which has the largest number of species, the dimension of its assembly and disassembly hypergraphs only allowed us to compute its assembly and disassembly holes up to dimension 2. Despite this technical limitation, we found that MDSine system is enriched with 2-dimensional disassembly holes (*p* = 0.005, Fig. 5j). Except from the disassembly skeleton of Stein system, for all other skeletons, low- and high-dimensional holes do not co-occur (colors in Fig. 4f-j). These results indicate an over-representation of assembly and disassembly holes for empirical systems.

**Figure 5:**
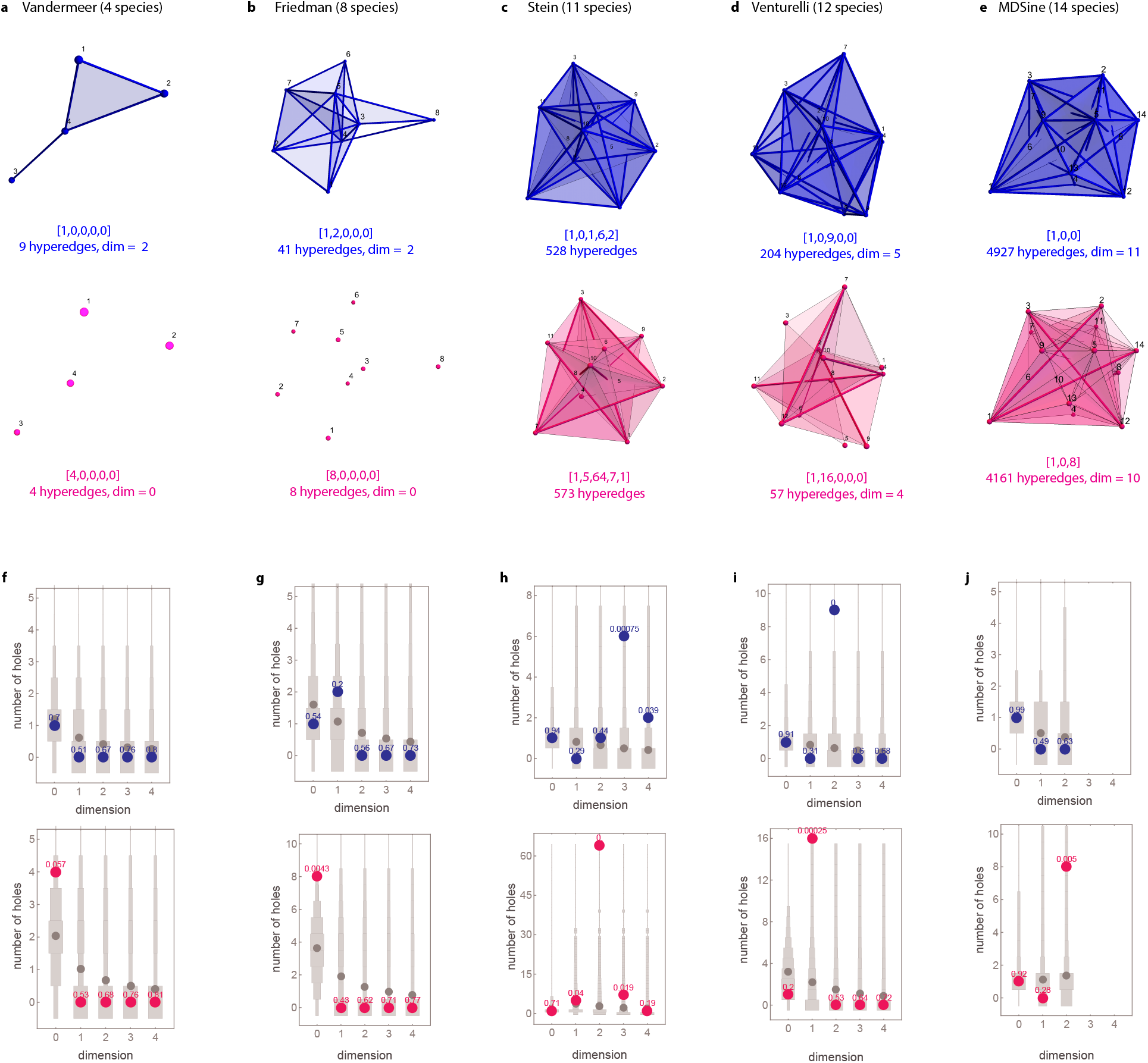
Empirical ecological systems are enriched with assembly and disassembly holes. **a. to e.** Embedding of the assembly (blue) and disassembly (pink) hypergraphs for the empirical interaction matrix and empirical intrinsic growth-rate vector. **f. to j.** Statistical analysis of the number of empirically observed assembly (blue) and disassembly (pink) holes with respect to the number of holes observed in the 3,000 randomization of the intrinsic growth-rate vectors (grey). Numbers above each color dot denotes the probability of observing those many holes among the 3,000 randomizations.

## Discussion

Ideally, assembly and disassembly hypergraphs should be entirely constructed by experimentally testing the coexistence of each of the different species collections (Supplementary Note S6). Such an experimental task is feasible for systems with a few species^29^, or for systems where massive automated co-culture experiments are possible^30^. We followed this approach to construct the assembly hypergraph of *S* = 5 core bacterial species in *Drosophila Melanogoster* gut microbiota from the in-vivo experimental data obtained by Gould et al29 (Supplementary Note S6.1). The assembly hypergraph we get has one connected component and two one-dimensional assembly holes, confirming the ubiquity of coexistence holes in ecological systems. However, for systems with a large number of species, the combinatorial explosion 2*^S^* – 1 in the number of different species collections makes it impossible to rely on experiments only. Circumventing this limitation requires parameterizing population dynamics models to predict the coexistence of some species collections. Our work uses the LV model, whose parameterization requires minimal data due to the simplicity of the model (Supplementary Note S3.4). While future work should not overlook the explanatory and prediction power of the LV model, we expect that coexistence holes become even more likely by using more complex models^31^.

Overall, our work shows that the structure of species coexistence is a complex interplay between filled spaces (i.e., species collections that coexist) and empty spaces (i.e., species collections that do not coexist). Some empty spaces give rise to assembly and disassembly holes, characterizing cases where coexistence abruptly breaks. Assembly holes represent unexpected obstructions to build larger species collections. Namely, they characterize collections that cannot coexist despite that their sub-collections can. Disassembly holes represent extremely “fragile” species collections that can be hard to preserve since removing any of its species will cause a collapse of many other species. Theoretically, we found that these coexistence holes are unavoidable, and we corroborate that these holes are more frequent than expected by chance across experimental systems. Importantly, the emergence of both classes of coexistence holes can be explained by simple dynamics under biotic and abiotic constraints, revealing the order shaping the assembly and disassembly of ecological systems. Note that classic coexistence studies in ecology^10,11^ are based on the assumption that coexistence holes do not exist, which can provide misleading results. Indeed, identifying discontinuities in ecological systems can improve our ability to understand which processes in nature are driven by internal constraints of design and not simply by randomness as it is more commonly believed.

## Acknowledgments

MTA gratefully acknowledges the financial support provided by CONACyT grant No. A1-S-13909. Funding to SS was provided by NSF grant No. DEB-2024349.

## Author contributions

MTA, CS, and SS designed the study. MTA, LM, and AK performed the theoretical analysis. MTA, CS, and SS wrote the manuscript.

## Data accessibility

The code supporting the results will be archived on Github.

## Methods

### An homology theory to uncover the homotopic skeleton of arbitrary hyper-graphs

A mature theory with efficient algorithms is already available to calculate the Betti numbers for the special class of hypergraphs known as *simplicial complexes*—hypergraphs where each hyperedge contains all its boundaries^32,33^. However, to the best of our knowledge, there is no efficient algorithm to calculate the homology (Betti numbers) for arbitrary hypergraphs. We note that hypergraphs are well-studied combinatorial objects^34^, with increasing applications accross the sciences^35–37^. Mathematically, the reason why the theory for simplicial complexes does not apply to arbitrary hypergraphs is because the topological space generated by hypergraph is neither open nor closed (Supplementary Note S2). Our approach circumvents this problem by using a new subdivision algorithm that transforms an arbitrary hypergraph into a simplicial complex (Definition 12 in Supplementary Note S2.2, and the example of Supplementary Fig. S3). The key property of our subdivision algorithm is that the resulting simplicial complex has the same homology as the original hypergraph (Theorem 1 in Supplementary Note S2.2). We further prove that the subdivision can be calculated by computing the Vietoris-Rips (VR) complex associated to a graph built from the hyperedges (Proposition 1 in Supplementary Note S2.3). This last result enabled us to compute the Betti numbers and hence the homology of hypergraphs using efficient algorithms developed for VR complexes^38^. An accompanying `Julia` package (with interface to `R`) provides all functionalities introduced in this paper.

### Constructing assembly hypergraphs from population dynamics

We take a structuralist approach^12,16^ to study how assembly and disassembly holes emerge in ecological systems. Specifically, we assume that the biotic conditions and governing laws of a system characterize the internal constraints defining the structure and dynamics of ecological systems, while abiotic conditions can change freely and characterize the potential diversity within such internal constraints^15,23^. By using this structuralist approach with our formalism, we can quantify the possible assembly and disassembly skeletons that can be observed in a given system (i.e., the “forms” that the ecological system can adopt) as the environment changes (i.e., as abiotic conditions change).

To make the above approach operative, we leverage on the mathematical tractability of the Lotka-Volterra (LV) population dynamics model^39^:

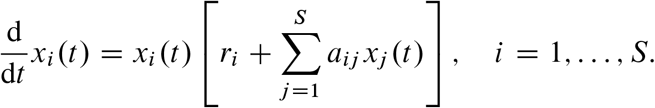

In this model, *x_i_*(*t*) represents the abundance of species *i* at time *t*. The parameters *r_i_* represents the intrinsic growth rate of species i, and *a_ij_* represents the effect that species *j* has on the percapita growth rate of species *i*. In the idealized governing laws described by the LV model, biotic conditions are phenomenologically captured by the 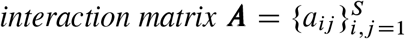, describing how species interact with each other. Abiotic conditions are phenomenologically captured by the 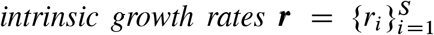, describing how species grow in isolation under given abiotic conditions. By using the LV formalism, we constructed the associated assembly hypergraph *H* by computing whether each of the 2*^S^* – 1 different species collections coexists or not under model parameters (***A***, ***r***). We defined coexistence following Jansen’s permanence criterion^40,41^ (Supplementary Note S3).

